# Remote cortical perturbation dynamically changes the network solutions to given tactile inputs in neocortical neurons

**DOI:** 10.1101/2021.06.10.447816

**Authors:** Leila Etemadi, Jonas M.D. Enander, Henrik Jörntell

**Affiliations:** Neural Basis of Sensorimotor Control, Department of Experimental Medical Science, Lund University, Lund, Sweden

## Abstract

The neocortex is a widely interconnected neuronal network. All such networks have a connectivity structure, which limits the possible combinations of neuronal activations across it. In this sense, the network can be said to contain solutions, i.e., for each given external input the cortex may yield a specific combination of neuronal activations/output. If the cortex has a variety of states, a given input could result in a range of possible outputs. There will also be a vast range of outputs that are not possible due to the network structure. Here we use intracellular recordings in SI neurons to show that remote intracortical electrical perturbation can impact such constraints on the responses to given tactile input patterns. Whereas each given tactile input pattern induced a wide set of preferred response states, when combined with cortical perturbation they induced response states that did not otherwise occur. The findings indicate that the physiological network structure can dynamically change as the state of any given cortical region changes, thereby enabling a very rich, multifactorial, perceptual capability.

## Introduction

Several studies have reported a wide distribution of cortical activity in response to input from a variety of modalities, directly or indirectly, indicating that the neocortex is in principle a globally interconnected network (Ferezou et al., 2007; Frostig et al., 2008; Fu et al., 2003; Ghazanfar and Schroeder, 2006; Hihara et al., 2015; Keller et al., 2012; Olcese et al., 2013; Rancz et al., 2015; Saleem et al., 2013). More surprisingly, information about the specific quality of tactile inputs is present in apparently any region of the neocortex (Enander and Jorntell, 2019; Enander et al., 2019; Genna et al., 2018), and thalamus (Wahlbom et al., 2021), and remote stroke-like lesions degrade the processing of tactile inputs in neurons in the primary somatosensory cortex (Wahlbom et al., 2019).

Without external input, the cortex has a complex, ever evolving internal activity (Hipp et al., 2012; Mantini et al., 2007; Stringer et al., 2019). This global cortical state can be defined as the distribution of activity across all neurons at any given point in time, and because of the sheer number of neurons the cortical state is extremely high-dimensional (Spanne and Jorntell, 2015; Stringer et al., 2019). Since the neurons are connected to each other, the network structure will constrain the number of possible states, or at least make some states, more or less, likely than others (Berkes et al., 2011; Golub et al., 2018; Luczak et al., 2009). This can be described as a system which potentially has an infinite number of input-output solutions, but which turns out to have a discontinuous landscape of solutions, i.e. where some solutions are much more probable than others, as in a state attractor (Ringach, 2009). That would mean that for a given tactile input pattern, for example, which is delivered randomly in relation to the current cortical state, there would be a tendency to form specific clusters of response types, or network solutions. This is indeed also possible to observe (Norrlid et al., 2021).

If the neocortex is sufficiently densely interconnected, an activity change in almost any individual neuron, or small group of neurons, would equal a change in the global cortical state, which in turn could impact the response to a given sensory input in any individual cortical neuron. Indeed, previous analyses have shown that perturbation of the activity even of single neurons will impact the responses of other nearby neurons (London et al., 2010), and intense single neuron activation can affect behavioral decisions (Houweling and Brecht, 2008; Voigt et al., 2008).

It should also be the case that perturbation of the activity of any neuron is theoretically possible to detect from any other single neuron regardless of its location in relation to the perturbation. The response of an individual neuron to a given input could be impacted even if the remote perturbation does not lead to any overt measurable response in it. This is because the perturbation could result in dynamic alteration of the global physiological network structure, i.e., the changes in the effective network structure that would result if some neurons dynamically fall below their activation threshold, which in turn could impact the network pathways that supply the neuron with inputs. Hence, an externally imposed perturbation could induce ‘new’ states in the cortex, i.e., states that does not otherwise arise, which should result in that the response states to a given input pattern deviate from the normal condition, at least temporarily, for as long as the effect of the perturbation lingers in the global cortical system. Clarifying such operational principles could give us important clues as to the structure of cortical processing and as well as for identifying limiting factors for successful communication in brain-machine interfaces. Here we explore the hypothesis that even remote temporary activity modifications could cause widespread dynamic changes in the effective cortical network structure by applying electrical, remote cortical perturbations while recording intracellularly from neurons in the SI cortex and providing specific tactile input patterns.

## Results

We recorded intracellularly from 19 putative pyramidal neurons between layers 2/3-5 (recording depths: 0.3 - 1.1 mm from the cortical surface) in the primary somatosensory cortex using the *in vivo* whole cell patch clamp technique (Fig. 1A). Because two of the neurons did not have an apparent response to tactile stimulation patterns, only 17 of these recordings were included in the main analysis, whereas the two nonresponsive neurons were included in one control test (see Fig. 4C). During recordings, we delivered a set of eight spatiotemporal tactile afferent activation (TA) patterns, or input patterns, each repeated in the order of 50-100 times (see Methods for details), in random order. While such tactile inputs can reach the SI neurons through direct cuneo-thalamo-cortical pathways, information about the tactile input is widely distributed in the cortex and could hence reach the recorded neuron through multiple parallel pathways (Fig. 1A). To test how interference with remote cortical networks could impact the responses in the SI neurons, we combined the TA inputs with remote cortical stimulation (CX). As illustrated in Fig. 1A, a localized CX perturbation may affect multiple cortical and thalamic network pathways, hence affecting their transmission state (‘Gates’). If such pathways would be partly responsible for meditating the tactile information to the recorded neuron, then there would for each given tactile input pattern be multiple alternative response types that could arise in the recorded neuron depending on the combined transmission states of all pathways impacting the neuron.

**Figure 1.**
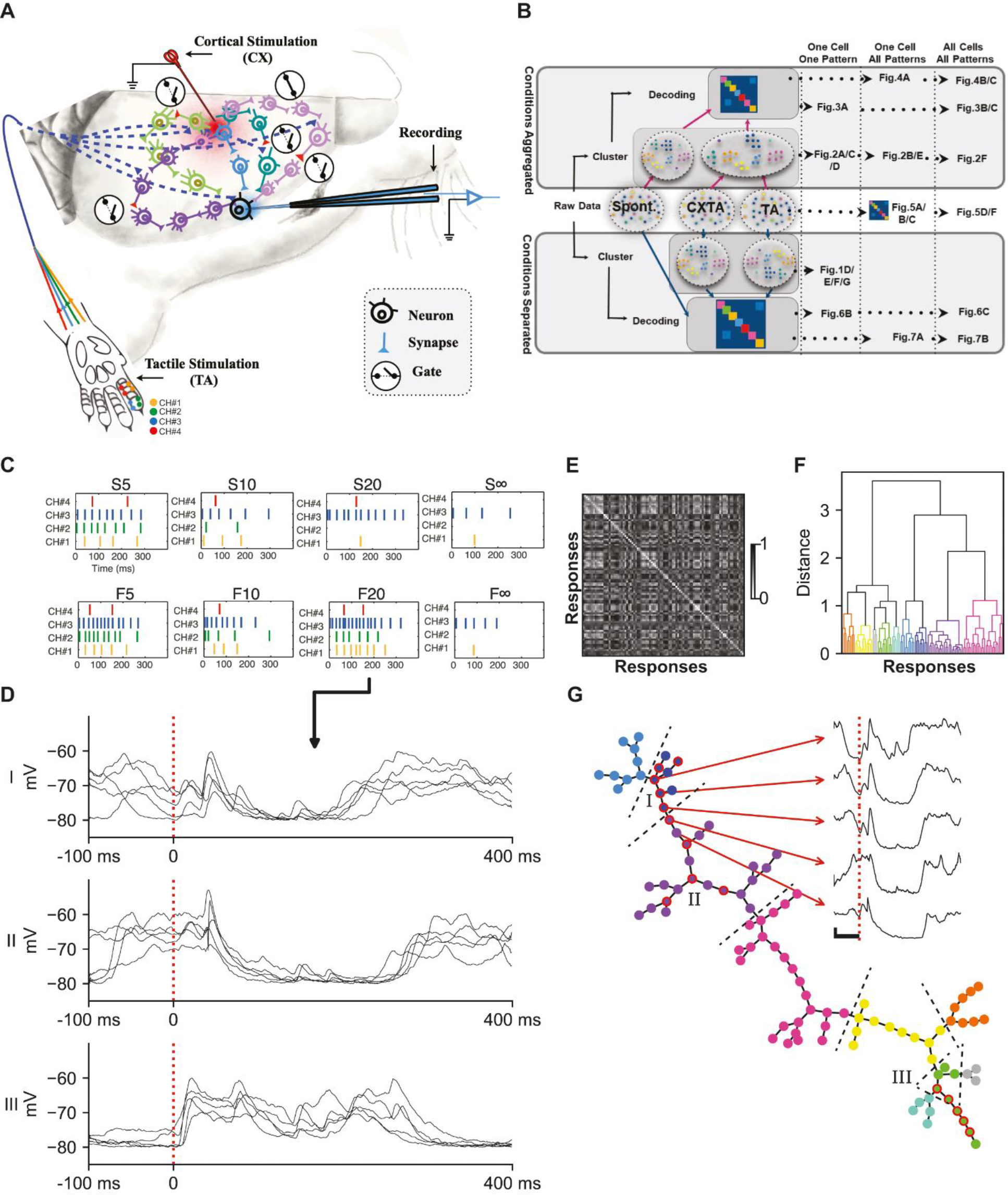
Experimental setup, analysis overview and illustration of the clustering method. **(A)** Experimental setup illustrated in schematic drawing. The second digit of the left forepaw was presented with eight spatiotemporal tactile afferent (TA) input patterns, delivered through an electro tactile interface distributed across four skin sites on digit 2 (channels, indicated with different colors) (bottom). Recordings were made in the SI cortex. The TA input patterns were delivered alone or conditioned by a cortical stimulation (CX) at a remote cortical site. In addition to the direct cuneo-thalamocortical input, the evoked tactile afferent activity distributes across the cortex through various neuronal network pathways, and the CX was intended to interfere with the transmission through these indirect routes (gating) before they reached the recorded neurons. **(B)** Overview of the analysis of the entire paper with keys to indicate which Figure shows which part of the analysis. The analysis was iterated for two different main hypotheses: (i) that the CX did not create unique clusters of responses for any given tactile input pattern and the responses evoked under the TA and CXTA conditions were therefore pooled before clustering (‘Conditions Aggregated’); (ii) that the CXTA and TA conditions evoked responses that were not part of the same distribution (‘Conditions Separated’). Note also that each analysis result could represent one cell and one input pattern, one cell across all input patterns, or all cells across all input patterns. **(C)** In the diagrams of the labelled TA input patterns, each vertical dash represents the timing of a stimulation pulse, and the color indicates through which channel it was delivered. **(D)** Example intracellular responses evoked by the same tactile input pattern (F20, the TA condition only), in one of the neurons. Each of the three panels show a subset of responses (N=5) from one cluster each (I-III). Onset of the TA input pattern is at time zero, marked with a dashed vertical red line, note that the duration of this specific pattern was 320 ms (as shown in panel C). **(E)** Distance matrix for all responses evoked by the TA input pattern F20. Note that the matrix is symmetric, and the upper triangle of the matrix is only included for visual clarity. **(F)** Dendrogram based on the distance matrix in (E) visualizing the cluster formation based on the estimated optimal cutoff (see Methods). Each response is colorized according to their designated cluster. **(G)** Proximity graph based on the dendrogram in (F). Each response (node) is connected to the closest response as defined by the calculated linkage matrix (see Methods). Each response is colorized according to its designated cluster (same colors as in F). Inset, five example responses spanning one cluster boundary, with the onset of stimulation indicated by a vertical dashed red line and calibration bars indicating 5 mV and 100 ms, respectively. The raw traces illustrated in (D) are indicated as nodes with red borders (i.e., subsets of clusters I-III). Dashed lines indicate boundaries between clusters.

**Figure 2.**
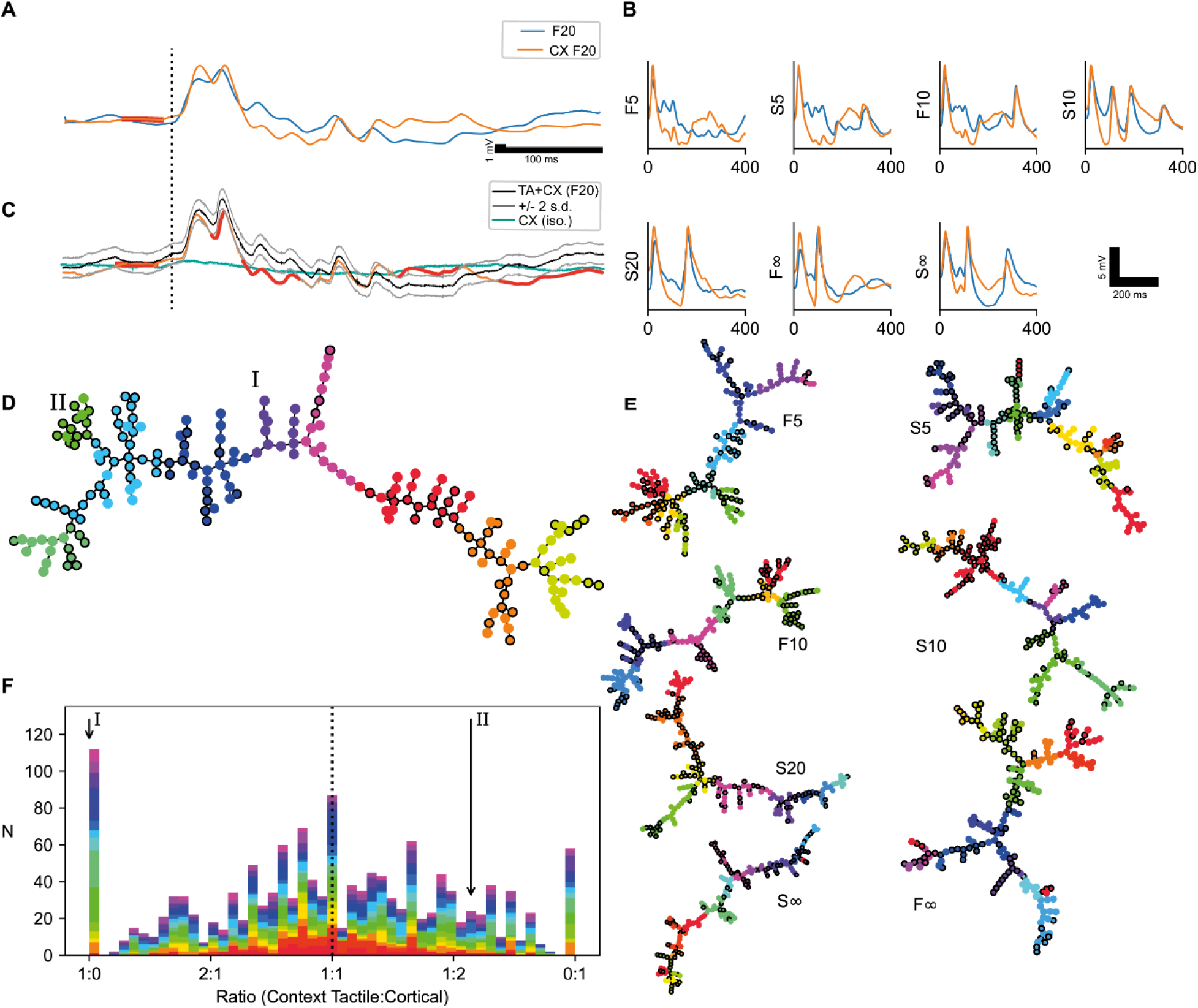
Impact of CX stimulation on TA responses and resulting sorting into response clusters. **(A)** Mean response to TA input pattern F20 with and without preceding cortical stimulation (CXF20 and F20, respectively). The onset of the TA input pattern is marked with a vertical dashed line. The CX stimulation occurred at the thick red horizontal line (stimulation artefacts blanked). **(B)** Mean responses for the other seven TA input patterns with the TA (blue) and CXTA (orange) conditions superimposed. In this case the pre-stimulus activity has been clipped. **(C)** Average CXTA response (orange) superimposed on the calculated algebraic sum of the TA and the CX responses (black). Thick red parts of the orange trace indicate where the CXTA response went outside the 95% confidence interval (the +/-2 standard deviation range (grey traces)) of the calculated response. Green trace indicates the CX response in isolation. In this panel only, non-smoothed responses are shown. **(D)** Proximity graph for the clusters identified for the aggregated F20 and CXF20 responses. The CXTA responses are indicated with black outlines. Two clusters have been marked; (I) is a cluster with only TA members; (II) is a cluster dominated by CXTA members (with a ratio of TA:CXTA responses of 1:3.6). **(E)** Corresponding proximity graphs for the seven other stimulation patterns of the example cell. **(F)** Histogram of the ratio of TA vs CXTA conditions for every single cluster from all recordings (the clusters from all 17 neurons for 8 stimulation conditions each). Each neuron is represented by a unique color. The locations of the clusters (I) and (II) from panel (D) are marked with arrows.

**Figure 3.**
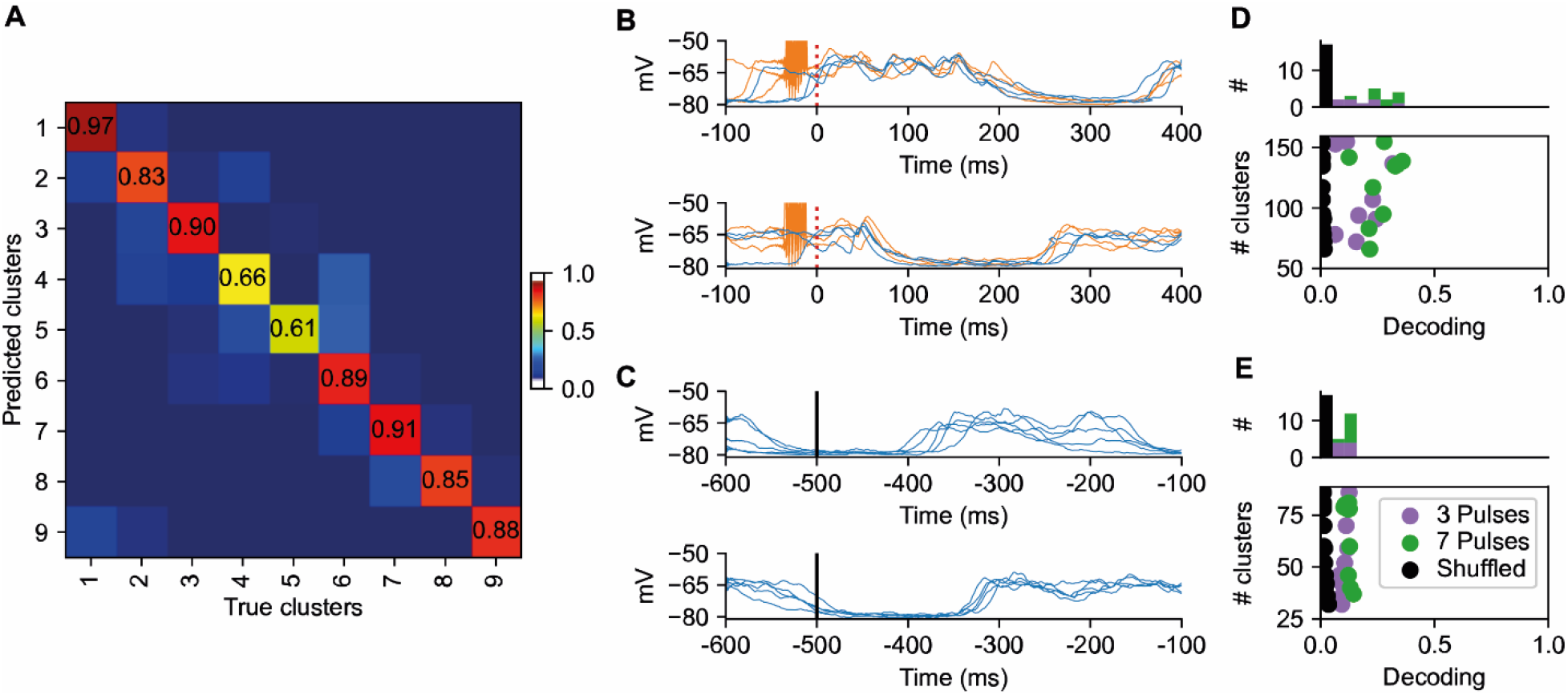
Separability of the response clusters. **(A)** Confusion matrix generated through PCA and subsequent kNN classification of clustered responses evoked by the F20 and the CXF20 input patterns (i.e., the TA and the CXTA conditions pooled) for the example neuron. The F1-score in this example was 0.85 with a chance level of 0.11 (1/9). **(B)** Superimposed raw traces from two example response clusters (Clusters 1 and 7). Note that both responses evoked under the TA condition (blue traces) and under the CXTA condition (orange responses) co-existed in the same clusters. The CX stimulation artefacts, that preceded the onset of the tactile input pattern (red dashed line), are clearly visible in the orange traces. **(C)** Examples of clusters identified in the spontaneous activity, which were recorded in the time window before the onset of the TA input pattern (− 500 to −100 ms). **(D)** The sum of the number of clusters for each neuron (N=17) across all eight patterns (TA and CXTA pooled), plotted against its average decoding performance (cluster separability). Neurons for which we used seven CX pulses are indicated in green, whereas neurons with three CX pulses are indicated in purple. Black dots indicate the controls, one for each neuron, where the cluster labels were shuffled. Note that the Y axis does not start at zero. **(E)** Similar display as in (D), but for clustered spontaneous activity.

**Figure 4.**
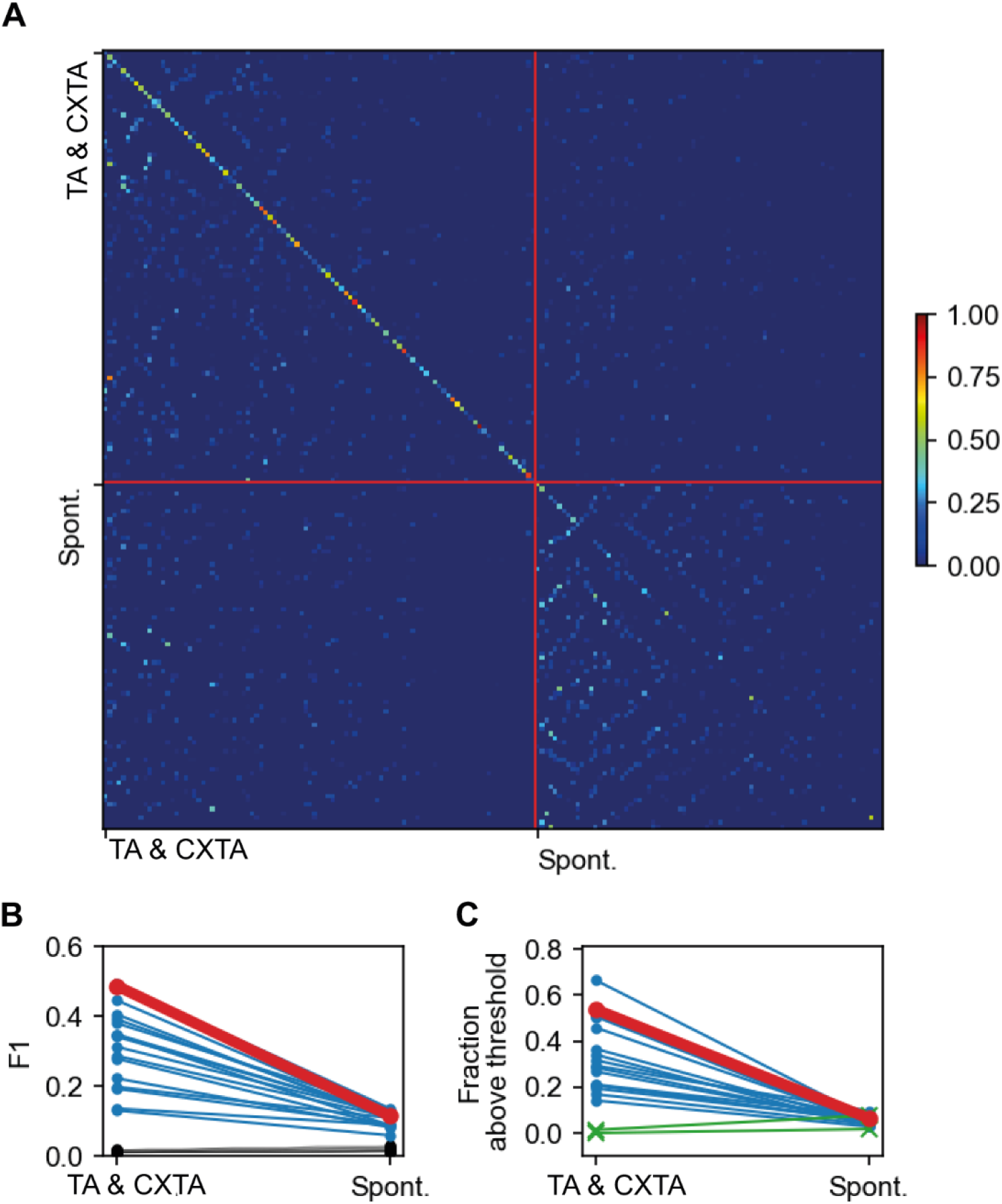
Separability of clusters calculated across all input patterns and spontaneous activity. **(A)** Separability of identified clusters when comparing all clusters from all input patterns in one recording (TA & CXTA Conditions Aggregated), as well as all clusters from spontaneous activity, for the example neuron. Red lines indicate boundaries between evoked and spontaneous responses. The F1 scores for the evoked responses were 0.49 (chance level=0.010; TA & CXTA compared to TA & CXTA) and for the spontaneous activity 0.12 (chance level=0.013; Spont. compared to Spont.), respectively. **(B)** Across all neurons recorded, the F1 scores of the evoked versus the spontaneous responses. The cell illustrated in A is indicated in red. Chance levels are shown in black. **(C)** Similar display as in A but instead reporting the fraction of clusters with an individual F1 score above a defined threshold. The threshold was defined as the weighted mean F1 score plus two times the weighted standard deviation calculated from the response clusters of the spontaneous activity for each recording. This panel also illustrates the 2/19 cells without any detectable response to TA inputs (green).

### Introduction to the cluster analysis

We used a cluster analysis to identify if alternative response types/response states could be evoked by the exact same stimulation pattern. The cluster analysis was performed on the responses (time-voltage curves) evoked by the TA input patterns and by the combined cortical and tactile, CXTA, input patterns, respectively. In order to establish whether clusters of responses were separable from each other, we used a decoding analysis. As shown in Fig 1B, this two-stage analysis was applied to test two different hypotheses, in turn. The first hypothesis was that CX did NOT impact the intracellular neuronal responses to a given TA input pattern, hence the responses evoked by the corresponding TA and CXTA stimulations were pooled (‘Aggregated’ in Fig. 1B). The results of the analysis of this hypothesis are presented in Figures 2-4. The second hypothesis tested was that the CXTA stimulation condition created a set of responses that were separated (‘Separated’ in Fig. 1B) from the TA responses, even though the TA input pattern component was identical between the two conditions. The results of the analysis of this hypothesis are presented in Figures 6-7. We also used the decoding analysis to test if the unclustered responses evoked by the TA and CXTA stimulation patterns were different from each other (Fig. 5). First, we introduce the clustering procedure by using the example of the responses evoked by one of the TA inputs patterns (Fig. 1D-G).

**Figure 5.**
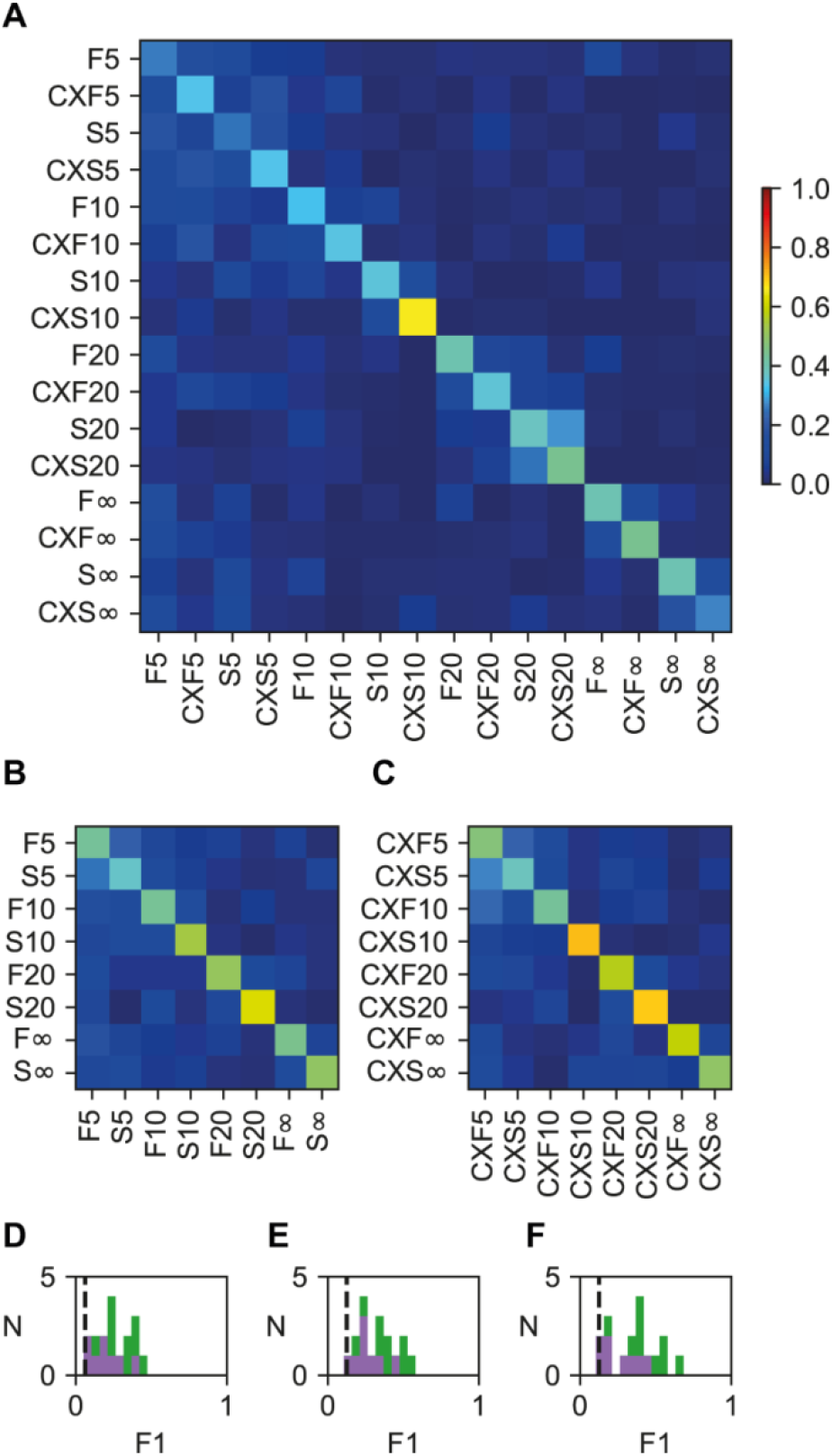
Separability of non-clustered responses. **(A)** Decoding of non-clustered TA and CXTA (separated) responses for the example cell with a F1 score of 0.38 (chance level=0.062, i.e., 1/16). **(B)** Decoding of non-clustered TA responses for the same cell (F1 score=0.49; chance level=0.125). **(C)** Decoding for non-clustered CXTA responses for the same cell (F1 score=0.55; chance level=0.125). **(D-F)** F1 scores for the population of recorded cells for non-clustered responses (green and purple bars indicate cells with seven and three CX pulses, respectively) for both conditions (D, F1=0.25±0.11; shuffled control=0.05±0.004), for the TA condition (E, F1=0.32±0.12; shuffled control=0.12±0.005), and for the CXTA condition (F, F1=0.36±0.15; shuffled control=0.12±0.006). Chance level is indicated by the vertical dashed black line.

**Figure 6.**
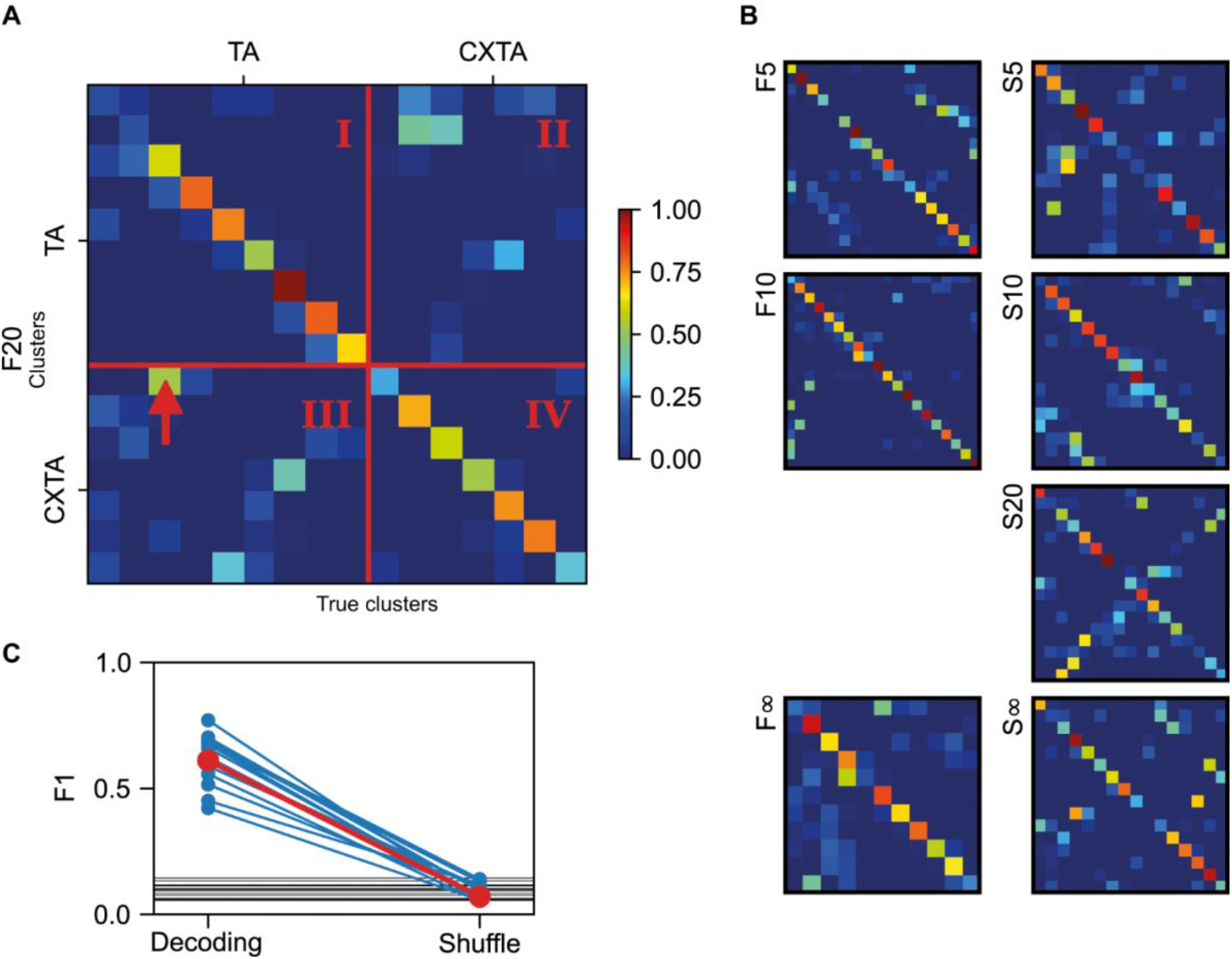
Separability of the response clusters evoked by the same input patterns but under different conditions. **(A)** Confusion matrix indicating the decoding of the response clusters to input pattern F20, where the clusters were formed separately for the TA and CXTA conditions (F20 and CXF20 ‘Separated’). The red lines indicate the boundaries between the different conditions (True:Predicted annotations are I=TA:TA, II=CXTA:TA, III=TA:CXTA, IV=CXTA:CXTA). The arrow points to an example element with confusion where the true condition (TA, according to the clustering) did not match the predicted condition (CXTA) as defined by the decoding analysis. **(B)** Confusion matrices similar to the one in (A) for the remaining seven tactile input patterns. **(C)** Average F1 scores across all neurons. The red line indicates the decoding level for all response types identified for each tactile input pattern, averaged across all eight input patterns, before and after shuffling of the response type labels, for the sample neuron shown in A and B. The blue lines indicate the average decoding level for the other 16 neurons. The horizontal black lines indicate the mean chance level per neuron.

**Figure 7.**
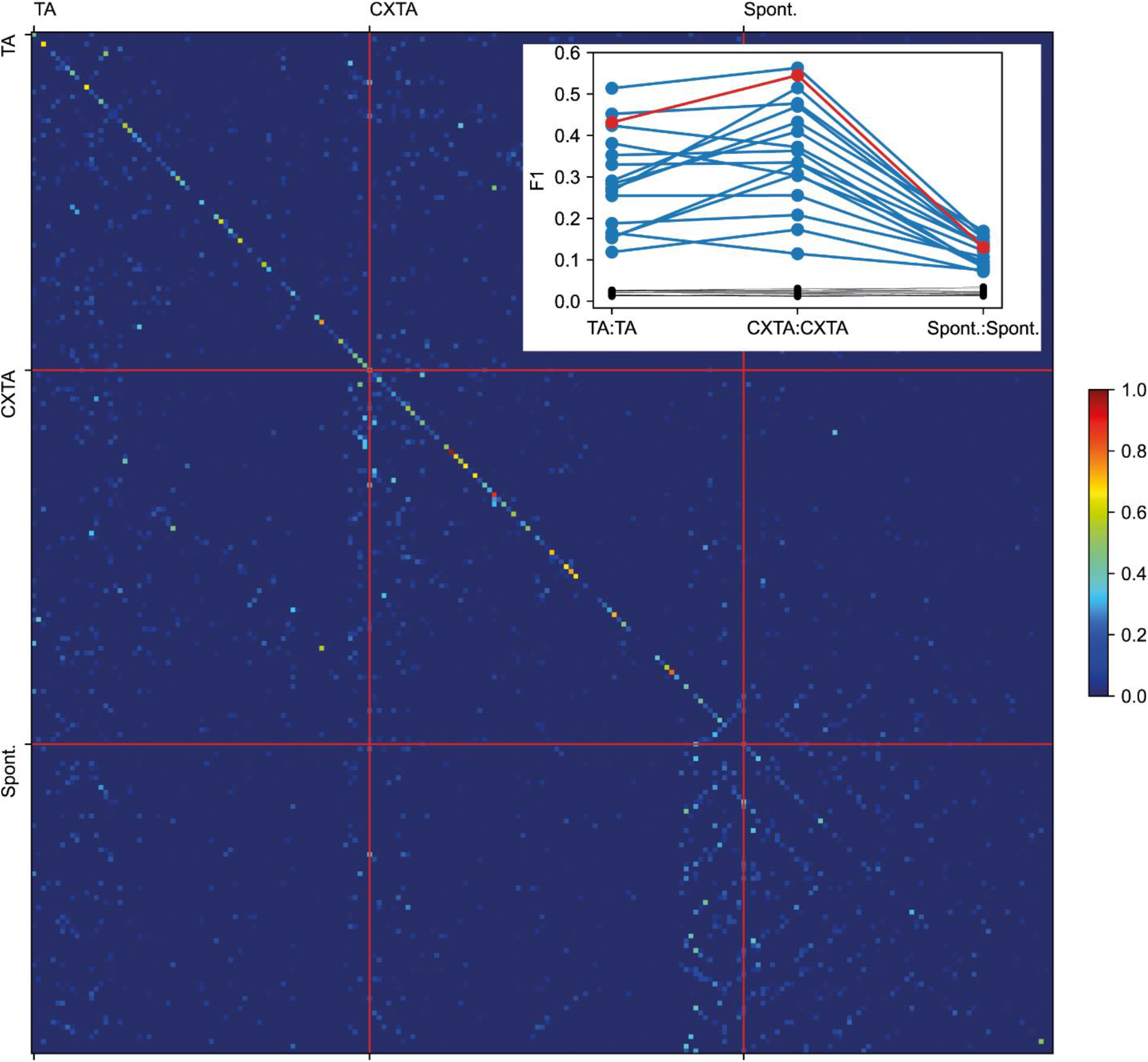
Cluster separability across all input patterns and conditions. The separability of identified clusters when compared to all clusters from all input patterns and both conditions (separated conditions as in Fig. 6), as well as all clusters from spontaneous activity, for the example neuron. Red lines indicate boundaries between the conditions. The F1 scores (chance level in parenthesis) were for TA:TA 0.43 (0.014); for CXTA:CXTA 0.55 (0.013); and for Spont.:Spont. 0.13 (0.015). Inset at top right shows the F1-scores under the separate conditions for each neuron. Black lines indicate the respective chance levels. The example recording is indicated with red line.

In order to limit the analysis to the synaptic inputs each neuron receives, the recordings were made with a mild hyperpolarizing bias current to minimize spiking activity. Hence, the intracellular voltage from a single neuron reflects the states in the subset of cortical neurons directly in contact with the recorded neuron. These states are affected by internal activity as well as by externally imposed inputs. Accordingly, even though the exact same input pattern (‘F20’, see Fig. 1C) was repeatedly delivered, the individual raw responses could differ substantially from each other (Fig. 1D). However, although the responses could vary extensively, there was also a tendency for some responses to be more similar to each other, suggesting that some preferred response states, or types, did recur (I, II, and III, respectively, in Fig. 1D). The clustering analysis quantified this tendency (Fig. 1E) and found that for all the responses it was a general rule that some responses were more closely related to each other than they were to the rest of the responses (Fig. 1F). As shown in Figure 1G, each cluster (or response type) had different numbers of ‘members’. ‘Adjacent’ responses could have a somewhat gradual change in appearance, but border transitions between two separate response types were characterized by the loss or addition of distinct response features (Fig. 1G, raw traces), suggestive of a dynamic gating-out or gating-in of specific subsets of the network pathways mediating the tactile input (Fig. 1A).

### Impact of combining the TA input with CX stimulation

When the F20 TA stimulation pattern was conditioned by a preceding CX stimulation (the CXTA condition, indicated as CF20 in Fig. 2A), the average response changed. Although there was some overlap in the initial components of the two mean responses (TA versus CXTA), later components diverged more clearly (Fig. 2A). Also, across the other seven TA patterns, the CXTA condition appeared to impact the average response (Fig. 2B). Notably, the CX stimulation on its own did not have any major response in the recorded neuron (Fig. 2C, green trace), which was to be expected for a remote stimulation. Moreover, we directly tested if the CXTA response deviated from the algebraic sum of the TA and the CX responses. We found that the average CXTA response fell outside the constructed TA+CX response by more than 2 standard deviations for more than 5% of the response time in all cases except one (for the remaining cases, we recorded a range of 8-77% of the time spent outside the +/− 2 s.d range; N=88 cases of comparisons, based on data from the 11 cells which included the CX stimulation, across 8 stimulation patterns). Hence, these findings were compatible with that the remotely located CX stimulation (Fig. 1A) primarily caused indirect interference with the subnetworks mediating the TA inputs to the recorded neurons (Fig. 1A).

### Cluster analysis for the two stimulation conditions ‘aggregated’

Our analysis first had the working hypothesis that there was no difference between the responses evoked by specific input patterns under the TA and CXTA conditions, i.e., whatever clusterability of the responses could be found it was not going to be due to the condition under which the response was evoked. Therefore, for each of the eight input patterns, we pooled responses evoked under the TA and the CXTA conditions, yielding a total of eight input patterns. Each individual input pattern was analyzed separately using the cluster analysis method, with the main purpose of exploring whether the hypothesis could be falsified. Figure 2D illustrates the clustering of the responses evoked by the F20 pattern, under both the TA and the CXTA conditions. A falsification of the hypothesis could be said to have been reached if some clusters consisted only of responses of the TA or of the CXTA condition, for example.

For one of the clusters (indicated by I in Fig. 2D) this criterion was reached. Although all other clusters in Fig. 2D did contain both conditions, the ratio between the two conditions varied substantially between clusters, where for example the cluster II was dominated by the CXTA condition. Figure 2E illustrates the same analysis but for the seven other input patterns, in which cases there were several clusters in which only responses evoked by one of the conditions were found. The cell had a total of 97 clusters (12.1 clusters in average per input pattern), of which 5 clusters contained TA responses only and 3 clusters CXTA responses only. Figure 2F summarizes the results of this cluster analysis for all stimulation patterns across all cells. From this analysis it is clear that the bulk of the clusters did contain both conditions, though with a ratio between the conditions that varied substantially, but there was a disproportionate number of ‘single condition clusters’ that would seem to falsify the hypothesis that the response clusters did not depend on the condition under which the response was evoked.

### Decoding analysis identifying the degree of separability of the ‘aggregated’ clusters

We next used a PCA+kNN decoding analysis to quantify the separability of the responses identified as belonging to specific clusters. If the observed clusters were not separable, but instead only arbitrary boundaries drawn across noise variations in the responses, the decoding analysis would help identifying that. In contrast, the result of this decoding analysis indicated that for the example neuron and the example input pattern F20, the separability, or the decoding accuracy, of the clusters was high (Fig. 3A). This can be seen by the high values of the cells in the diagonal of the matrix, whereas the number of misclassified responses, non-zero values outside this diagonal, was low. The average F1 score, which we used to quantify this separability, was 0.85 in Fig. 3A. Fig. 3B illustrates raw data from two of the response types/clusters/states in Fig. 3A, where it can be seen that response states under the TA and the CXTA conditions could indeed in some cases be highly similar.

### Clustering analysis of the spontaneous activity

In order to explore to what extent recurring random fluctuations could cause the clustering observed, we also applied the clustering algorithm to the spontaneous activity and found that the algorithm identified response clusters also in the spontaneous activity (Fig. 3C). We are therefore next compared the separability of the clusters of the evoked responses versus those of the spontaneous activity. Fig. 3D illustrates the average F1 score for the evoked responses (i.e., the average F1 score across the eight confusion matrices, such as the one illustrated in Fig. 3A, illustrated for each cell). The number of pulses used in the CX condition (3 or 7, illustrated in different colors in Fig. 3D) did not significantly affect how many clusters were observed (Mann Whitney U, U-statistic=31.5, p-value=0.35). It can be noted that the number of clusters identified for each cell could vary quite a lot, but this did not appear to impact the separability of the clusters (Linear regression N=17, Slope=58.9 (CI95%=−81.6 - 199.4) R^2^=3.1%, p=0.5). Fig. 3D also illustrates the shuffled control decoding for each cell, where the cluster ID# were shuffled between the responses evoked by the same input pattern before the PCA+kNN analysis was reapplied. Since the decoding accuracy collapsed following the shuffling, the identified clusters were confirmed to be separable by the PCA+kNN analysis across the population of recorded neurons. Figure 3E illustrates a similar analysis for the spontaneous activity, and found that the PCA+kNN decoding results were clearly above chance and that separability disappeared when the cluster#-ID of the spontaneous responses were shuffled. This result indicates that also the spontaneous activity contained separable clusters. This was not surprising, since it merely indicates that the spontaneous activity does not follow fully random time evolutions, but instead abide by certain rules that are most likely due to the structure of the cortical network. The fact that the spontaneous activity is not random has been extensively explored before (Berkes et al., 2011; Luczak et al., 2009). Hence, our clustering algorithm was powerful enough to find clusters even when there was no peripheral input. However, the separability of the clusters of spontaneous activity was substantially lower (F1 score 0.11+/−0.016, Fig. 3E) than for the clusters formed among the evoked responses (F1 score 0.22+/−0.09 Fig. 3D).

### Decoding analysis for the ‘aggregated’ clusters identified across all TA patterns

We next applied the decoding analysis across all the clusters across all eight TA patterns per neuron. In this analysis, we also included the clusters identified for the spontaneous activity in the same neuron, to verify that the clusters of the spontaneous activity were not the same as the clusters of the evoked responses. As shown in the resulting giant matrix (Fig. 4A), it did indeed turn out that the responses evoked by the different TA input patterns formed clusters that were separable from each other. This was remarkable, considering the very high number of clusters involved in this analysis, and hence a very high risk of confusion. However, there were a few exceptions where some response types (clusters) were not well separated from the other clusters (i.e., dark blue squares in the diagonal of the matrix of Fig. 4A).

Interestingly, the clusters generated for the spontaneous activity were as a rule not well separable. But then it should be noted that the spontaneous activity was first split up depending on which TA input pattern it preceded and then clustered separately – the confusion matrix then compares spontaneous clusters preceding specific TA input patterns. Since the spontaneous activity could not predict which TA input pattern that was about to be delivered it was not surprising that these clusters were confused with each other. Conversely, the contribution to the limited above-chance decoding of the spontaneous activity almost exclusively arose from the clusters associated with the first of the input patterns that went into the classifier (i.e., the upper left 1/8 within the spontaneous clusters in Fig. 4A). And in line with these observations, confusion between the different clusters (elements outside the central diagonal in the matrix) for the spontaneous activity mostly arose with other clusters of spontaneous activity. Hence, this part of the analysis verified that when responses were similar to each other, they would also be confused and thereby collapsing the F1 score reported by our analysis.

The F1 score for the elements of the matrix belonging to the evoked responses was much higher than the F1 score for the spontaneous clusters, which is shown in red for the illustrated cell in Fig. 4B. This was also the rule for all the neurons (Fig. 4B). Figure 4C instead reports how many clusters (as a fraction of all clusters of the same stimulation pattern) that had an F1 score above a threshold detection level based on the separability of the spontaneous activity clusters. In this plot, we also included the 2/19 cells that did not have a clearly defined response to the TA inputs. Unlike all other neurons, these 2 neurons fell down to a value close to zero in this analysis and thus served as a control that neurons without meaningful clusters could be singled out by the PCA+kNN verification method. Hence, we conclude that the PCA+kNN could be used to indicate that the identified clusters were indeed separable in the 17/19 neurons included in the analysis.

### Decoding analysis of un-clustered responses

Hence, from Figs 2-4, we conclude that the hypothesis that the TA and the CXTA responses were part of the same pool of responses was contradicted by the cluster analysis. Before fully testing the alternative hypothesis, that the response types evoked under the TA and the CXTA conditions were separated from each other, we first tested this idea without a preceding cluster analysis. Hence, all the raw responses of the 16 stimulation conditions were compared to each other using the same type of PCA+kNN decoding analysis (Fig. 5) as performed in the confusion matrices above. Figure 5A illustrates for one neuron that the un-clustered responses evoked by the different input patterns were indeed separable from each other, though not perfectly so; the F1 score was in this case 38% (compared to a chance level of 6.2%, 1/16). Interestingly, there were several cases where the differences between the TA responses of different input patterns (F5 and S5 for example) could be exceeded or matched by differences between the responses evoked by same tactile input pattern under the TA and CXTA conditions, respectively (F5 and CXF5, for example). Moreover, when comparing the eight input patterns for each condition separately (Fig. 5B for TA; Fig. 5C for CXTA) the F1 score did not rise substantially compared to Fig. 5A (55% and 49%, respectively, compared to a 12.5% chance level). Figure 5D-F summarizes the result of this analysis for all neurons recorded. Overall, these results were compatible with that the differences between the responses evoked under the TA and CXTA conditions for the SAME TA input pattern could be comparable to the differences between the responses evoked by different TA input patterns under the same condition. This suggested that the clusters of TA and CXTA responses could be separate from each other.

### Decoding analysis of clustered responses with conditions ‘separated’

Therefore, we next performed the clustering analysis with the hypothesis that the TA and CXTA conditions represented separate pools of responses. For each given tactile input pattern we clustered the responses for the TA condition separately from the responses for the CXTA condition (Fig. 1B, lower half). The diagonal of the sample confusion matrix (Fig. 6A) did indeed indicate a high degree of specificity of the response clusters generated separately for the two conditions. However, this was not a rule without exceptions. Note for example the square indicated by the red arrow in Fig. 6A, which indicated that this particular set of responses had a high risk of confusion between the two conditions (i.e., responses generated under the CXTA condition were relatively often misclassified as belonging to the third cluster of the responses evoked under the TA condition). Across the seven other stimulation patterns for this cell (Fig. 6B), the risk of confusion appeared to vary substantially, with the highest confusion risk being observed for responses evoked by the S20 pattern. On the other hand, for example the F10 pattern seemed to generate responses that were consistently correctly classified to the respective clusters, with much less confusion between conditions. Fig. 6C summarizes the results of this analysis for all cells, with the results from the eight different input patterns being averaged for each cell. Here it can be noted that overall, this analysis provided support for the hypothesis that the responses evoked by the TA and CXTA conditions did really represent distinct sets of responses, even though the tactile component of the input pattern was exactly same.

### Decoding analysis for the ‘separated’ clusters identified across all TA patterns

Similar to Figure 4, but now under the hypothesis that the responses evoked under the two conditions were separate sets of clusters, we also compared all the responses recorded under all TA patterns (Fig. 7). These results showed in addition to the results of Fig. 6 that also when the decoding analysis was performed across all clusters of one example neuron, the response clusters across all stimulations were confirmed to be largely distinct and well above the F1 score for the clusters of the spontaneous activity (in the example neuron, the F1 score (chance level) for TA:TA was 0.43 (0.014), for CXTA:CXTA 0.55 (0.013), and for Spont:Spont it was 0.13 (0.015)). Across the population of neurons (inset in Fig. 7), this was a consistent pattern. Interestingly, the clusters of the CXTA condition were more distinctly separable than those of the TA condition. This could be due to that whereas under the TA condition the tactile input patterns combined with an uncontrolled spontaneous activity, the CX stimulation may have constrained this spontaneous activity to a somewhat tighter range, hence making the CXTA responses more separable.

## Discussion

We found that remote perturbation of cortical activity profoundly impacted the intracellular responses in SI neurons to a set of fixed spatiotemporal patterns of tactile afferent input. We showed that even though each given input normally generated a range of response states, the remote cortical perturbation could induce new types of response states to the same tactile input. Our results show that the cortical network contains a wide range of network solutions for any given input and that the specific solution that apply at a given moment likely depends on an interplay between the activity levels across wide areas of the cortex.

We have previously reported that the intracellular responses evoked in a neuron by the exact same tactile input pattern differ from one repetition to another and that these responses tend to sort into separate preferred response types, which are unique for each neuron (Norrlid et al., 2021). Here we extended that analysis by showing that a cortical perturbation can alter the preferred response types/states to any given tactile input pattern. Our cluster analysis indicated that when the hypothesis was that TA and CXTA responses were not different, several clusters would be occupied only by members of one of the conditions (Fig. 2F). In a response decoding analysis without clustering, Fig. 5 provided support for a separability of the responses evoked under the two conditions (TA and CXTA) using the same tactile inputs. Figures 6 and 7 further showed that for the responses evoked by the same tactile input, the clusters identified for each stimulus condition separately were indeed separable.

Despite the observed induction of new network solutions, i.e., input-output relationships, our remote cortical perturbation did not generate distinct synaptic responses in the cortical neurons (Fig. 2C). We showed that the combination of remote cortical stimulation and the tactile input pattern (CXTA) produced a response that was significantly different from the algebraic sum of the CX and TA stimulations. Hence, the remote cortical perturbation caused its main effect by interfering with pathways that mediated the tactile afferent input to the recorded SI neurons, via cortical pathways (Fig. 1A) naturally with possible involvement of subcortical structures such as the cuneate nucleus and thalamus.

The intracellular signal reflects the activity of the ~10000 neurons connected to the recorded neuron. These afferent neurons could in principle have had completely independent activity in which case the recorded signals would have been noise plus the contribution from the TA input. However, instead of noise, our analysis indicated a discontinuity in both the evoked and the spontaneous intracellular responses, which therefore displayed multiple preferred response states. This is a logical consequence of that the neurons are connected to each other, directly and indirectly, since such connectivity constrains the possible response combinations across the neuron population, in line with the observations from previous recordings of population level neuron activity (Berkes et al., 2011; Golub et al., 2018; Luczak et al., 2009). It is important to point out that the fact that the cluster analysis identified multiple specific response states does not imply that these response states are fixed entities or the only response states possible. The cluster analysis merely indicates that there are some central response features that some responses share, whereas at least one of these features were not shared by the members of the other clusters. Superimposed on these features were more fine grained differences between responses within clusters (Figs 1 & 3), which with the present dataset size and clustering method could not be further separated. The several different response states observed for each input, even as we explored input merely from distal digit 2 combined with CX, indicate that the cortex is potentially capable of an extremely high number of solutions.

We found a discontinuous landscape of solutions also for the spontaneous activity. According to the decoding analysis, they were less separable than the response states evoked by any of the TA inputs. This can be explained by that the skin stimulation tended to induce more specific subsets of states in the network. Each of the eight TA input patterns created a relatively unique subset of response states, indicating that the effective network structure/cortical state did adapt to the external input to some extent.

### Methodological issues

It is likely that the anesthesia limited the range of possible spontaneous activity patterns. We do not think that this invalidates the main observed phenomenon that the cortical perturbation tended to shift the state of the integrated network. This notion comes from an analysis of the multiple intracellular response states induced by the same TA input patterns used here (Norrlid et al., 2021), where we found that the response states arose both during the synchronized ECoG state, the probability of which is increased by the anesthesia, and during desynchronized activity, which is more similar to the predominant state observed in awake animals (Constantinople and Bruno, 2011). Anesthesia could for example increase the transmission of the perturbing signal as a consequence of the lowered general activity present in desynchronized states. But the fact remains that even early anatomical estimates indicate that any neuronal signal can reach any other neuron within just a few synapses (Arbib et al., 1998) – since then, more advanced neuroanatomical techniques have revealed that the extensiveness and promiscuity of cortical neuronal axon branching have been widely underestimated in the previous literature (Gerfen et al., 2018) (https://www.janelia.org/project-team/mouselight; https://mouse.braindatacenter.cn/). Hence, even though the control of the gating of these pathways in the awake animal may be better organized, the potential contributions of the communication revealed by our findings will not go away when the animal is awake.

Our cortical stimulation does not reflect normal activation patterns within the brain and may hence push the global network into temporary states it would otherwise not enter. This is in fact the case for any method using any form of active interference with the cortical activity, including Brain Machine Interfaces. We do not however think that this potential problem prevents us from drawing the conclusions that we make. The principle that any variation in cortical activity, for example due to changes in ‘thoughts’ or external cues from the same or from other modalities, will impact the response to a given sensory input, should anyway be valid, though the magnitude of the impact may be more subtle than what we observed in our experiments.

### Relationship to the literature

There are several previous studies that show that the internal state of the cortex will affect the responses to sensory inputs (Arieli et al., 1996; Curto et al., 2009; Fiser et al., 2004; Hasenstaub et al., 2007). A difference with the present study is that we had a more highly resolvable set of inputs, and in addition a highly resolved temporal response evolution across several 100 ms (Norrlid et al., 2021). Another difference is that we did not define the cortical state by any arbitrary measure but instead let the data indicate the presence and number of response states that could arise, whereby the resolution of our state analysis could also be higher. We have previously shown that the global ECoG state (synchronized versus desynchronized) does not have a predictive value for which response state will be induced by the TA input patterns (Norrlid et al., 2021). This may be due to that each afferent neuron is connected to specific sets of neurons, and most likely the response state that will be induced depend on the state in the specific subnetworks those afferent neurons are located in, where each individual afferent subnetwork may not necessarily follow the global ECoG state at all times. Hence, each neuron ‘sees’ unique circuitry components of the global state, which can also be shown by that each neuron generates responses that are complementary to the responses of other neurons to the same input (Oddo et al., 2017), and each of these components may behave somewhat differently, which would create the response states we observed, and which was the reason for the proposal of the concept of the ‘multi-structure’ cortical state (Norrlid et al., 2021). We believe that these multi-structure cortical states are dynamic phenomena, that would risk to be overlooked following any form of response averaging (for a recent example, see Diamanti et al. (2021)), and were hence so far not well accounted for in the literature.

Our results provide insights into how perceptual processes in the cortex can work at the neuronal circuitry level. The combination of a tactile input pattern with the neuronal response state, which can be widely different across neurons processing the exact same tactile input (Norrlid et al., 2021), will define the activity distribution across all neurons of the cortical circuitry – which is a potential definition of a ‘percept’. Our results indicate that the neuronal response states can be made even more diversified by specific activation patterns in remote cortical areas. Hence, any internal state variation, potentially a ‘thought’, or any external cue, no matter how small, can impact the activity distribution resulting from any given tactile input pattern, which provides for an extremely rich, high-dimensional perceptual capability. The principle of perceptual constancy, i.e. a constant percept despite differences in the underlying sensory input pattern (Garrigan and Kellman, 2008) (see also Purves et al. (2011)), requires that the perceptual decision criteria are robust to a wide range of resulting states, i.e. that multiple network solutions are perceived equivalently. How that process is coordinated mechanistically is a crucial issue for future studies, but it is clear from the present study that the mechanisms involved likely involve extremely rich dynamics engaging large parts of the cortex.

### Experimental Procedures

#### Ethical approval

The ethical approval for this study was received from the Lund/Malmö local animal ethics committee in advance (permit ID M13193-2017).

#### Surgical procedures

We made acute *in vivo* recordings in male Sprague Dawley rats (N = 15; weight = 250 – 380 g). Animals were initially prepared in the same way as in a previous study (Norrlid et al., 2021), briefly according to the following procedure: 1) The animal was sedated by inhaling air mixed with isoflurane gas (3%, for ~ 2 min); 2) To induce general anesthesia, a mixture of ketamine/xylazine (ketamine: 40 mg/kg and xylazine: 4 mg/kg, accordingly) was injected intraperitoneally; 3) an incision in the inguinal area of the hindlimb was made to insert a catheter in the femoral vein for continuous infusion of Ringer acetate and glucose mixed with anesthetic (ketamine and xylazine in a 20:1 ratio, delivered at a rate of ~5 mg/kg/h ketamine). After the initial preparation steps, the somatosensory cortex (SI); was exposed by removing a small part of the skull on the right-hand side (~2 x 2 mm), located at (from bregma): Ap: - 1.0 - +0.1, ML: 3.0-5.0. Also, another exposure (about the same size) was created in the skull to place the cortical stimulation electrode (CX), from bregma: Ap: −3.1, ML: 2.5, where the tip of the stimulation electrode was placed at the estimated depth of layer 5. Furthermore, a surface electrocorticography (ECoG) electrode was placed in the rostral part of the second cortical exposure.

The anesthetics used were chosen because they have previously been reported to not dramatically alter the neuronal recruitment order in spontaneous activity fluctuations and in stimulation-evoked responses as compared to the awake animal (Luczak and Bartho, 2012). As we have previously discussed extensively (Norrlid et al., 2021), anesthesia was required in order to achieve identical tactile stimulation patterns (where the electro tactile interface was the key, but would not be accepted in the awake animal) over a sufficiently long period of time. It also served to minimize brain activity noise caused by uncontrollable movements and internal thought processes unrelated to the stimuli. The level of anesthesia was assessed both by regularly verifying the absence of withdrawal reflexes to noxious pinch of the hind paw and by continuously monitoring the irregular presence of sleep spindles mixed with epochs of more desynchronised activity, a characteristic of sleep (Niedermeyer and da Silva, 2005). In order to prevent the exposed areas of the cortex from dehydrating, and to decrease the brain tissue movements, a thin layer of agarose (1%) was put on the exposed cortical areas. The animal was sacrificed by an overdose of pentobarbital at the end of the experiment.

#### *In vivo* neural recordings

Intracellular recordings were made in the whole-cell current-clamp mode. The patch-clamp pipettes were pulled to impedances of 6 – 10 MΩ from borosilicate glass capillaries using a Sutter Instruments P-97 horizontal puller. The pipettes were back-filled with an electrolyte solution containing (in mM): potassium gluconate (135), HEPES (10), KCl (6.0), Mg-ATP (2), EGTA (10). The solution was titrated to 7.35–7.40 using 1 M KOH. The signal was amplified by an EPC-800 patch-clamp amplifier (HEKA Elektronik) in the current-clamp mode (bandwidth from DC up to 100 kHz). The data was digitized at 100 kHz using the CED 1401 mk2 hardware and recorded using the Spike2 software (Cambridge Electronic Devices, CED, Cambridge, UK).

In the next step, the patch pipette was inserted into the SI neocortex (Paxinos and Watson, 2006), first in a stepwise approach and with applied positive pressure to the electrode tip, in order to penetrate through the dura and the pia without blocking the electrode. The electrode was subsequently advanced slowly (~ 0.28 – 0.3 um/sec) through the cortical tissue until it encountered a neuronal spike, evoked by electrical skin stimulation to digit 2 (single pulse stimulation using 1-4 channels (see below), intervals between stimulations 0.3 s). Once the spike signal magnitude dramatically increased, the positive pressure was switched to a brief negative pressure and a mild hyperpolarizing current (in the order of −10 pA) was applied to the electrode to facilitate the formation of a GigaOhm seal. The access to the intracellular space was then obtained by brief, episodical negative pressure applied to the electrode tip. Once the electrode had proper access to the intracellular environment and showed a stable signal, the data recording began. The data included in the analysis are from neurons with stable <−55 mV membrane potential in down states, with a peak-to-peak between up and down states of >10 mV and a spike amplitude of >25 mV before and after starting the protocol. All neurons recorded were putative pyramidal neurons rather than interneurons based on that they exhibited infrequent bursts of two or three spikes but had an absence of longer bursts or sustained periods of high firing (Luczak et al., 2009). Note that for each electrode track made in the cortex, before gaining intracellular access we also recorded the local field potentials (LFP) evoked by the electrical tactile stimulation, to identify the minimal latency time of arrival of the synaptic cortical responses from the skin stimulation. All recordings were made in layer II/III-V (at depths of 350 – 1100 um).

#### Electrical tactile and cortical stimulations

The experiments consisted of delivering tactile stimulation patterns through an electro tactile interface, consisting of bipolar pairs of percutaneous stainless steel needle electrodes (isolated except for the tip (0.2-0.5 mm) and inserted into the superficial part of the skin of the second digit of the contralateral (left) forepaw), and combining them with intracortical electrical perturbations. The skin had four such bipolar pairs of electrodes, each pair being a channel of tactile input (Fig. 1A). The bipolar skin stimulation electrodes were delivering stimuli where each pulse was a 400 μA constant current pulse, with a duration of 200 μsec, which is about two times the threshold for activating tactile afferents using this type of stimulation (Bengtsson et al., 2013), and below the threshold required for activating nociceptive afferents (Ekerot et al., 1987).

The electro tactile interface was used to deliver spatiotemporal tactile afferent (TA) stimulation patterns. We used eight different TA stimulation patterns (0.5fa, 0.5sa, 1.0fa, 1.0sa, 2.0fa, 2.0sa, flat fa, flat sa; Fig. 1C) that were delivered in a preset randomized order (the same protocol as in Enander et al. (2019); (Norrlid et al., 2021); Oddo et al. (2017); Wahlbom et al. (2019)). Each TA stimulation pattern lasted for less than 350 ms, and consecutive TA patterns were separated by a randomized interval of about 1.8 s (Oddo et al., 2017).

Unlike our previous experiments, the present set of experiments included cortical perturbation, an electrical CX stimulation with a remote location relative to the recordings (the distance between the CX electrode and the recording electrode was ~ 7 mm) made in the SI cortex (medial ‘parietal’ cortex, Fig. 1A) that could be combined with the TA input patterns. A glass-insulated tungsten electrode (exposed tip 100-150 um, (Jorntell and Ekerot, 1999)) served as the CX micro-stimulation electrode, with its reference ground inserted into the neck muscle. The CX stimulation consisted of 400 μA constant current pulses and 0.2 ms pulse duration, consecutive pulses were separated by 3 ms, and we used either 3 or 7 pulses in different experiments. The current intensity and the number of pulses were selected so as to avoid that the CX stimulation evoked any detectable response in the recorded neurons in the SI cortex. A ‘CXTA’ stimulation consisted of the CX stimulation combined with a TA stimulation, where the CX stimulation was timed so that the last pulse of cortical stimulation preceded the first TA stimulation pulse by 17 ms. There were 8 TA stimulation patterns (see above) plus 8 CXTA stimulation patterns, where the CXTA patterns consisted of the fixed CX stimulation combined with each of the 8 TA patterns.

We used a premade stimulation protocol consisting of the TA patterns and CXTA patterns, i.e., 16 patterns of stimulation, mixed in random order. Each protocol consisted of 50 repetitions of each pattern i.e. 800 stimulus deliveries. For neuron recordings that lasted longer than the duration of a full protocol, we first made a recording of the responses to the CX stimulation in isolation (100 repetitions, at 1 s intervals), and then restarted the premade protocol for another 50 repetitions/800 stimulus deliveries. The material included 19 neurons, where two neurons were discarded from the main part of the analysis, due to a lack of a clearly detectable response to the TA inputs. Of the remaining 17 neurons, N=6 neurons lasted for two full repetition sets (N=100 repetitions) of the premade protocol, i.e., they hence received 1600 stimulus deliveries (800 TA inputs and 800 CXTA inputs). For the remaining neurons, we obtained recordings for 30, 31, 46, 49, 50, 50, 50, 68, 70, 90 and 94 repetitions of all TA/CXTA patterns. We obtained recordings of CX stimulation in isolation, 100 repetitions, for 11 of these neurons. Note that we have previously shown that even larger number of repetitions of the tactile afferent input patterns than we used here does not lead to detectable plastic differences in the responses they evoke in SI neurons (Wahlbom et al., 2019).

#### Post processing: response filtering

In order to facilitate the post processing analysis of the responses (Fig. 1B), we first created post-processing responses from the raw data. In order to define a stable baseline voltage, the recordings of the neuronal membrane voltages were linearly detrended in 10 second long segments. The recordings were subsequently low pass filtered using a simple moving average with a width of 1 ms. In the recordings, where occasional spikes appeared, the onset of these spikes were found using a spike shape template and a recursive fitting algorithm (Mogensen et al., 2019). The spikes could subsequently be removed from the recording data by subtracting the mean signal of the 50 nearest surrounding occurences of the spike. Finally, the recordings were downsampled to 1 kHz.

#### Definition of evoked responses

The time window included in this analysis started 5 ms after the onset of the TA stimulation (i.e., for both TA and CXTA stimulation patterns) and ended 400 ms later (some TA patterns lasted almost 350 ms, and responses could sometimes be detected at least 50 ms after the last stimulation pulse). The 5 ms gap relative to the TA onset corresponded to the conduction time from the skin stimulation to the earliest possible arrival of synaptic responses in the neocortex, as judged by the LFPs.

#### Definition of analysed spontaneous activity

Analysis of spontaneous activity was based on the recorded neuron activity during the −500 - −100 ms time window relative to the onset of each TA stimulation (defined as time 0).

#### Clustering method

We used clustering analysis in order to identify the different response states that could be evoked on different repetitions of the same stimulation pattern (for both CXTA and TA patterns, ‘Aggregated’ or ‘Segregated’, see Fig. 1B).

The purpose of the clustering algorithm was to detect distinct groups of response states, where the response state equaled the time-voltage curve of the evoked response. Evoked response states that were more similar to each other than they were to all other responses evoked by the same stimulation pattern were considered to belong to a cluster, i.e., the same group of responses. The purpose of the clustering algorithm was hence to identify groups where the ‘members’ of each group had to be internally similar, while also being distinct relative to other responses. Furthermore, the algorithm had to be able to automatically determine the number of clusters that could exist, since this could not be known *a priori*.

Each response was z-score normalized, i.e., the response mean over the whole time period was subtracted from each response and then the remainder of each response was divided by the standard deviation, and then a distance matrix was calculated for the whole set of responses. The distance matrix was populated by calculating the M number of Principal Components (PC) explaining 95% of the variance for N-1 responses. All responses were then fitted to the PCs using the least square method. The resulting PC coefficients were used to position the responses in the M-dimensional PC space. Within this M-dimensional space, the Euclidean distances between a reference response and the rest of responses were calculated. The distances between the reference response and each of the other responses were then put in a matrix (the distance matrix; Fig. 1E), and the procedure was repeated until each response had been the reference. Finally, the distances were normalized.

From the Euclidean distance calculations above, a linkage matrix (Fig. 1F) was calculated using the hierarchical clustering approach of Ward’s minimum variance method (Ward, 1963). This method sequentially finds the two closest neighboring responses with respect to their Euclidean distance from each other, and their Euclidean distance is indicated in a dendrogram such as in Fig. 1F (this ‘cophenetic distance’ is indicated as the distance along the y axis). Then it finds the pair of the second closest neighboring responses and so on until there are no responses left. However, a response may also be closer to a center point between a pair of responses than it is to other responses, in which case the cophenetic distance to that center point is assigned to the response. Sometimes, the shortest distance that can be found is between two such center points. All these distances are used to create the hierarchical structure of responses, where the distances between all such pairs can be compared (the distance along the y axis of Fig. 1F).

While following the above principles, there were some special situations that could arise. A cluster was assumed to be valid only if it had five or more members. All non-valid clusters were put in the same “undefined” cluster. In addition, for one of the cells the situation arose that one of the clusters consisted of only one member. In this case this response and cluster was excluded from all subsequent analysis and displays.

The next issue is to identify the cophenetic distance at which the identified clusters are objectively the most separable. The y axis in Fig. 1F (the full range of which corresponds to the largest total cophenetic distance) was then divided into 100 steps. For each level of cophenetic distance, we calculated the number of clusters found for the actual data. Data shuffling was achieved by shuffling the position of each response along the X axis of the dendrogram (Fig. 1F), and therefor disrupting the relationship between cluster and response.

We next calculated the separability of the clusters by using a decoding analysis (described in detail below). The decoding analysis yielded, for each clustering result, a measure of the separability of the clusters as the F1-score. With the F1-score, we could apply Gap statistics to identify the cophenetic distance providing the largest cluster separation relatively to the shuffled data. For each of the 100 cophenetic distances considered, we obtained a F1 score for the non-shuffled data and another F1 score for the shuffled data. Since the specific value of the chosen cophenetic distance defines the number of clusters to be considered, we could identify continuous stretches of cophenetic distances where the number of clusters remained unchanged. Over each of these continuous stretches we calculated a mean F1-score, and subtracted the mean F1-score of the shuffled data from the actual mean F1-score, which yielded the Gap statistic (Tibshirani et al., 2001). The middle value of cophenetic distance of the stretch with the maximum Gap statistic was used to define the cophenetic distance at which the clustering was objectively the most distinct. The clustering obtained at this value was then used for the remainder of the analysis and displays. The identified clusters using this approach are indicated by color codes in Fig. 1F for an example cell and example TA stimulation pattern.

#### Visualization of hierarchical proximity

Linkage matrices are commonly visualized as dendrograms (Fig. 1F). They display hierarchical data in a 2D-manner, and the relationship between clusters can be easily seen. However, they fail to display the detailed relationships between the constituent responses of the clusters. For example, the gradual but distinct transformation of one cluster into another proximal cluster (inset traces in Fig. 1G) cannot be seen using a dendrogram. Therefore, we also used an additional visualization technique.

The linkage matrix was transformed into a topological ‘Proximity’ graph by iterating through each defined bifurcation of the dendrogram (Fig.1F). Each bifurcation was considered to describe an edge and the branches describing the nodes connected by this edge. If a branch did not terminate in a specific response but instead into a center point between responses/clusters, then the edge was connected to the response within that cluster that had the least distance to the reference response on the other side of the bifurcation. The weight of the edge was set to be equal to the distance – 1, since they are conceptual inversions of each other meaning that a long distance is the same as having a low weight edge connecting them. The layout of the resulting proximity graph (Fig. 1G) was finally optimized using the Kamada-Kawai algorithm (Kamada and Kawai, 1989).

#### Evaluation of cluster separability using decoding analysis

A general decoding algorithm to determine the specificity of the response clusters was used throughout this article. It is the same decoding algorithm used by our lab in several previous publications (Enander and Jorntell, 2019; Enander et al., 2019; Wahlbom et al., 2019; Wahlbom et al., 2021), including intracellular recording data with multiple response states to each tactile input pattern (Norrlid et al., 2021).

The responses and their respective label were split into a training- and a test set stratified by cluster label frequency. The training set was used to train a PCA model explaining 95% of the variance, and the coefficients obtained by fitting the responses with the least-square-method to the principal components were calculated for both the training set and the test set. Finally, a kNN-classification was performed with the training set coefficients as training data, and the test set coefficients as test data. This decoding algorithm was performed 30 times, each time with a new split of train and test data. The F1-score was calculated from the average classification result across the 30 repetitions.

The decoding analysis was also repeated with the randomly shuffled cluster labels, reported as the “shuffled” context. This was done to establish the actual chance level of the data, as opposed to the theoretical chance decoding level (which equals 1 divided by the number of clusters). Finally, the shuffling was repeated three times to obtain the mean F1-score for the shuffled context.

## Author Contributions

LE and HJ designed the experiments. LE performed the experiments. JMDE designed and performed the clustering analysis. LE, JMDE and HJ analyzed the results, and wrote the article.

## Acknowledgements

This work was supported by the EU H2020 Grant FETOpen project# 829186, ‘ph-coding’ (Predictive Haptic COding Devices In Next Generation interfaces)

